# Genes near tRNAs are enriched in translational machinery

**DOI:** 10.64898/2026.03.12.711363

**Authors:** Caroline West, Lauren Dineen, Abigail Leavitt LaBella

## Abstract

Transfer RNAs (tRNAs) are known for delivering amino acids to the growing polypeptide chain during translation. They can also influence gene expression, especially in times of nutrient starvation, through differential tRNA expression and modification. tRNAs have a highly consistent cloverleaf structure, but relatively few known regulatory elements govern this conserved structure despite the 20 different standard isotypes. This study examines gene enrichment patterns near tRNA in 1154 fungal genomes. Genes enriched in proteasome regulation, ion transport, and rRNA were found to be significantly closer to tRNAs than other pathways. These results were consistent across KEGG over-representation analysis (ORA), KEGG Gene Set Enrichment Analysis (GSEA), and Gene Ontology (GO) analysis. Proteasome, ion transport, and RNA are all important aspects of protein production and regulation, suggesting that genes required for the synthesis and quality control of proteins, including tRNAs, are located near each other. Protein regulation is an energetically expensive process, and local co-regulation could increase efficiency and stress impacts on proteins.

## Introduction

Outside of their role in translation, tRNAs have also been reported to modulate gene expression, such as tRNA abundance affecting translation efficiency, and uncharged tRNAs binding to regulatory elements to initiate or stall translation (Raina and Ibba 2014). tRNA binding to cytochrome c has been shown to inhibit apoptosis (Raina and Ibba 2014). tRNA can also transfer arginine to Arginyl-tRNA transferase (ATE1) to mark proteins for degradation by the ATP-dependent ubiquitin system (Ciechanover et al. 1985; Abeywansha et al. 2023). Furthermore, during stress response, tRNA can be cleaved into catalytically active tRNA fragments that can repress translation (Magee and Rigoutsos 2020; Yu et al. 2021). Given the myriad important roles of tRNA, dysregulation of tRNAs has been linked to cancer and neurodegenerative disease in humans (Orellana et al. 2022).

tRNAs are one of the most modified RNAs and go through multiple processing steps from transcription to maturity. When transcription is initiated for a tRNA gene, transcription factor TFIIIC binds to the internal promoters, the A box, and the B box (Talyzina et al. 2023). This then recruits a second transcription factor TFIIIB, which will bind upstream of the transcription start site (Talyzina et al. 2023). Next, polymerase III is recruited to initiate transcription, and this process can be repeated several times until it reaches a poly-T sequence signaling transcription termination (K C et al. 2024). This creates pre-tRNA, which contains a 5’ leader, potentially an intron, and a 3’ trailer. These must be spliced out, and CCA added to the 5’ end for later amino acid attachment (Li et al. 2022). Then the tRNA can become a mature tRNA capable of being charged with an amino acid and ready for translation (Chen et al. 2024). All 64 tRNA isoacceptors go through this process, and while they carry different amino acids, they consistently exhibit a highly similar cloverleaf structure. The variation primarily occurs in the variable arm and the anticodon stem loop, whereas the other components remain constant (Weiss et al. 2024). These known tRNA transcription mechanisms are unlikely to account for all observed changes in tRNA expression.

We hypothesize that genes near tRNA are enriched in translational machinery, which could influence tRNA regulation. To answer this question, we leveraged the y1000+ dataset, which includes 1154 genomes from the Saccharomycotina subphylum (hereafter referred to as yeast) and their functional annotations. Gene clusters are a known phenomenon in yeast where genes supporting similar functions colocalize together (Cera et al. 2019), but to our knowledge, large-scale genomics analysis of genes near tRNA has not been examined across a diverse eukaryotic lineage. We hypothesized that protein-coding and non-coding rRNA gene proximity could influence tRNA expression. We tested this by examining KEGG enrichment and GO enrichment in the genes near tRNAs.

## Materials and Methods

### Yeast Genomes

We obtained 1,154 yeast genomes and annotations from the y1000+ project (https://y1000plus.wei.wisc.edu/). This included functional annotations from the KEGG (Kanehisa 2000; Kanehisa et al. 2025) and GO (Ashburner et al. 2000; The Gene Ontology Consortium et al. 2023) databases. tRNA annotations were previously generated using tRNAscan-SE (Chan et al. 2021).

### Chromosome assembly visualization

To examine differences in tRNA and rRNA patterns across contigs, 26 of the 1154 yeast genomes had contig numbers of 16 or less that were equal to or within 1 of the literature reported number of chromosomes (Table S1). These genomes were plotted in R v4.4.2 (R Core Team 2024) with predicted tRNAs from tRNAscanSE and rRNAs predicted from Infernal v1.1.5 (Nawrocki and Eddy 2013). The tree illustrating the relationships between the species utilized the ape v5.8-1 package (Paradis et al. 2004) from R and was visualized using iTOL v6 (Letunic and Bork 2024).

### Gene distance calculations

To identify the protein-coding genes near tRNAs, we compared GFF genome files and tRNA files with bedtools v2.28.0 (Quinlan and Hall 2010). The “closest” function was run to populate a list of genes that were closest to every tRNA in every species and report the distance. Next, the genome files and tRNA scan files were run with bedtools “window” function set to 2000bp, which finds all the genes within 2000 base pairs of tRNA.

We subset the closest genes based on the tRNA’s isotype and codon preference. Relative Synonymous Codon Usage (RSCU) was used to classify codons into genome-specific preference categories. RSCU values were obtained from previous work (Zavala et al. 2024). tRNAs that decode codons with an RSCU less than 0.5 were “unpreferred”, RSCU greater than 1.5 were “preferred”, and RSCU 0.5 - 1.5 were “moderate. Any distances of −1 were excluded because this is bedtools way of reporting that there is no closest gene.

### Over-Representation Analysis (ORA)

To test which genes were enriched near tRNAs, we conducted Over-Representation Analysis (ORA). ORA takes an input gene list and compares it to a background gene list, to see which genes, functions, or pathways are statistically over-represented in the input list. Enrichment in KEGG annotations was conducted with the R package ClusterProfiler v4.16.0 (Yu et al. 2012) using the enrichKEGG function. Cluster profiler was run separately on subsets of the near-tRNA genes, including closest-only, the 2,000 bp window, tRNA isotype, preferred codons, unpreferred codons, and moderate codons. Cluster profiler has the following parameters: gene list of interest, universe or the background, keyType such as KEGG, organism, and p-value adjustment method. Organism was set to “ko” for KEGG Ontology to allow for comparisons that included multiple species. The method of multiple testing correction was Benjamini-Hochberg with an adjusted p-value cutoff of less than 0.05. The universe was all KEGG orthologs (KOs) possible in the 1154 yeast species.

For species-specific ORA, the organism code was “sce” for *Saccharomyces cerevisiae* and “yli” for *Yarrowia lipolytica*. The backgrounds were limited to KOs possible for those organisms.

### Predicting rRNA locations from Infernal

KEGG annotations do not include ribosomal RNA (rRNA). Therefore, we used Infernal v1.1.5 (Nawrocki and Eddy 2013) to identify rRNA genes in all the Saccharomycotina. Specifically, we used 8 CPUs per task and output the results in table format. The tRNAs were removed, and the outputs were converted to BED format for downstream analysis. Additionally, rRNAs annotated as bacterial or archaeal were removed. The Infernal documentation notes that there can be overlap in the models for bacteria, archaea, and eukaryotes in genes with similar homology across domains. Infernal was re-run with a few test genomes with the --nohmmonly flag and resulted in fewer bacterial or archaea results, but it had the same level of accuracy as the previous infernal run that had the bacteria and archaea hits manually removed afterwards. Re-running all of the genomes with Infernal --nohmmonly would be computationally expensive, and given that the results were very similar to the original results, the decision was made to keep the original results with the bacteria and archaea manually removed. Barrnap v0.9 (Torsten Seemann) was also run to compare to the original method because it is easier to select for just eukaryotes. Barrnap was substantially faster but was not as sensitive as Infernal.

### KEGG Pathways near tRNA genes

To further explore the ORA results, we found the distance from all members of specific KEGG pathways to tRNAs. KEGG Ontology (KO) IDs identified genes associated with the following pathways: Aminoacyl tRNA biosynthesis (map00970), glycolysis (map00010), fatty acid biosynthesis (map00061), ribosomal proteins (br01610, map03010), ribosomal pathway (map03008), tRNA modifications, and proteasome (map03050). Infernal v1.1.5 (Nawrocki and Eddy 2013) identification of rRNA was used to compare KEGG pathway identification of rRNA genes to rRNA gene prediction using a different identification process. For each pathway group, bedtools closest was run to calculate how far all the KOs in the group were from the closest tRNAs. The results were plotted in R as density plots using the package ggplot2 version 3.5.2 (Wickham 2016).

Statistical significance of differences in group distances was measured using a paired Wilcoxon test using R because the data were not normally distributed. The data was tested for normality using random sampling of 500 data points for each group to perform a Shapiro-Wilk test. Lack of normality was also confirmed with the Anderson-Darling test with the entire dataset from - 2000 base pairs to 2000 base pairs and with qq plots (Figure S1 and Table S2). Statistical significance of how far each group’s mean was from 0 was measured using a Wilcoxon signed-rank test in R.

### Gene Ontology (GO)

We used Gene Ontology to test enrichment for genes near tRNA. Gene Ontology (GO) analysis was done with the R Package topGO v2.60.1 (Alexa et al. 2006) using the elimination method. Only GO annotations with at least 10 core genes and a p-value of less than 0.05 were considered. This was performed on all the genomes in the Saccharomycotina separately and then combined to look for the top results across species. The results were plotted by enriched GO categories enriched GO terms were color coded based on category.

### Gene Set Enrichment Analysis (GSEA)

We used Gene Set Enrichment Analysis GSEA to test if ranking genes based on KO count yielded similar results to overrepresentation analysis. Gene Set Enrichment Analysis (GSEA) was run with the R package Cluster Profiler v4.16.0 (Yu et al. 2012) using the gseKEGG function and ranked by KEGG count.

### Visualizing tRNA locations at the chromosome level

*Yarrowia lipolytica* and *Saccharomyces cerevisiae* were chosen for chromosome level analysis due to their assemblies having the same number of contigs as expected number of chromosomes. Additionally, these species have experimental data from Oxford Nanopore sequencing on tRNA counts (*in prep*). From this data, we identified active and inactive tRNA genes using the sequencing read count data. Active tRNA genes were identified if they fell in the upper 95th percentile of the distribution of counts. Therefore, any tRNA gene with more than 11 reads was determined to be active.

The bedtools output for these species was read into R for ribosomal pathway KOs, ribosome protein KOs, tRNA scan, and rRNAs predicted by Infernal. Each of the components was set as a track with the location determined by the axis position. Horizontal width was determined based on the start and end points of the genomic element. All components were then combined with ggplot2 version 3.5.2 based on their chromosome.

## Results and Discussion

### Chromosome level assemblies reveal tRNA and rRNA distribution is variable across species

Yeasts are a highly diverse group of organisms, and there are considerable variations, such as chromosome number variation ranging from 4 to 16. In general, we observed that in most species (21 out of 26), rRNAs were found on 1-2 of their chromosomes. Conversely, some species (e.g., *Yarrowia lipolytica* and *Arxula adeninovorans* from the order Dipodascales) had rRNAs on every one of their chromosomes. Dipodascales species had a higher percentage of rRNAs on their chromosomes (ranging from 33% to 100%) than the other orders.

Saccharomycetales all had 25% or less of their chromosomes with rRNA. These findings illustrate the species-specific differences even within a subphylum.

In order to further investigate tRNA distributions across the 26 genomes, we visualized the genomes in R, plotting the different genomic elements as tracks. The genomic elements examined were tRNA location, ribosomal pathway genes predicted from KEGG, rRNA proteins from KEGG, and rRNA from Infernal. rRNA is a critical part of translation, so we wanted to see various aspects of its distribution and how that related to tRNA location. For *Yarrowia lipolytica,* we also had experimental data available for tRNA copy number. Previous studies have found a relationship between tRNA copy number and tRNA abundance (Bénitière et al. 2025), which could reflect tRNA availability and translational efficiency.

Despite having 510 predicted tRNAs, *Yarrowia lipolytica* only had 63 active tRNAs, and *Saccharomyces cerevisiae* had 275 predicted tRNAs with only 53 active tRNAs. The active tRNA count ranged from 12 to 34361, with an average of 828 in *Yarrowia lipolytica* and 2655 in *Saccharomyces cerevisiae*. In the *Yarrowia lipolytica* zoomed in view, 2 out of the 4 most highly expressed tRNAs are found near ribosomal pathway genes. rRNA and tRNA are both important for translation, so genomic organization of these two elements being in close proximity will be further examined. The zoomed in view of tRNA and other genetic elements can provide a snapshot into genomic organization, but the following sections use the entirety of the y1000+ dataset to uncover organizational trends across the 1154 yeast genomes and further investigates proximity of rRNA to tRNAs.

### Genes near tRNA are enriched in functional categories related to ribosomes and proteins

We hypothesize that gene content may influence the genomic organization of tRNA genes. To test this, we identified protein coding genes close to tRNA genes in 1,154 Saccharomycotina genomes. We identified 9,542 protein coding genes as the nearest gene to 199,965 tRNA genes. Bedtools cannot find the closest genes between contigs, so assemblies with a larger number of contigs return fewer results. A total of 2 tRNA genes did not have any protein coding genes nearby due to small contig sizes, and 10,231 did not return a closest gene. Bedtools does not report why a closest gene was not found, but it could also be attributed to contig differences. We also identified 10,527 protein coding genes within a 2,000bp window of the tRNAs.

We conducted KEGG over-representation analysis (ORA) on the 3,435 unique KEGG Orthologies (KOs) associated with the genes closest to tRNAs (Figure 3A.) This data was also subset into each individual tRNA isotype (20) and three levels of codon preference (Figure 3A). In total, we identified 16 pathways that passed multiple testing correction within each subset.

Across the dataset, we identified two types of pathways that were broadly shared: ribosomal and the neurological disease pathways (Figure 3A.) The two ribosomal pathways, “Ribosome” and “Ribosome biogenesis in eukaryotes” showed enrichment in 5 of the 9 total subgroups (Figure 3A). Ribosomes and tRNAs are both essential components of translation. The “Ribosome” pathway included the KOs such as K03685 (2 out of 9 subgroups), which is the ribonuclease III (RNT1 in *S. cerevisiae*), and K17440, the large subunit ribosomal protein (MRP49 in *S. cerevisiae*). The “Ribosome biogenesis in eukaryotes” pathway included KOs such as K11130 (NOP10 in *S. cerevisiae*, which is H/ACA ribonucleoprotein complex subunit 3), K02989, the small subunit ribosomal protein S58 (RPS5 in *S. cerevisiae*). The finding that tRNAs are enriched near ribosomal genes suggests genomic organization includes proximity of the key components of translation spatially near each other. There is also evidence that ribosomal proteins may play a direct role in tRNA gene expression; for example, ribosomal protein L11 regulates tRNA expression in human cell lines (Dai et al. 2010). Moreover, findings in *S. cerevisiae* suggest that RNA polymerase II modifies chromatin at both rRNA and tRNA genes (Yague-Sanz 2024). Having tRNA and rRNA genes close together genomically could allow for more efficient protein production, especially when having both elements in the same open chromatin region. This could reduce lag time if the organism is in need of rapid protein production or be an efficient way to conserve resources if protein production is low.

Multiple neurodegenerative human diseases were associated in the KEGG ORA analysis. Despite not having neurons, yeast have many of the cellular components and pathways that impact neurodegenerative disorders (Franssens et al. 2013). Generally, many of these neurodegenerative diseases have pathology related to protein misfolding and accumulation of protein aggregates. Several examples of proteasome regulation showed up in the ORA, including 26S proteasome regulatory subunit N3 (K03033), 26S proteasome subunit N2 (K03032), 20S proteasome subunit alpha 7 (K02727), and 20S proteasome subunit alpha 3 (K02728). Examples of proteasome dysfunction are Parkinson’s disease, where damaged proteins are allowed to accumulate, which leads to Lewy body formation (Mbefo et al. 2015), expansion of polyglutamine regions, which occurs in Huntington’s disease (Zhao et al. 2018), and amyloid proteins, such as with Alzheimer’s Disease (Kryndushkin et al. 2013). Mutations in these proteins or gene regions often lead to protein aggregation, leading to disease progression (Koszła and Sołek 2024). Homologs of these disease-associated proteins also occur in yeast cells and produce similar phenotypes when altered. This makes yeast a useful model for these diseases (Ocampo and Barrientos 2011) despite being generally single-celled. The two top KO categories associated with the neurodegenerative were proteosome (229 out of 743) and transport (320 out of 743, Table S3.) For example, K02725 (PSMA1) and K02726 (PSMA2) both have annotations for 20S proteasome subunits. We also found enrichment in K03936, which is an NADH dehydrogenase (ALI1 in *C. albicans)* and K02272 (COX8 in *S. cerevisiae)* for cytochrome C, both of which assist with electron transfer. Cytochrome C has also been found to bind to tRNA, and its binding affinity is dependent on the redox state of cytochrome C (Liu et al. 2016). Strong binding of cytochrome C has been shown to prevent apoptosis (Liu et al. 2016). Additionally, previous work has linked import of tRNAs into the mitochondria with the ubiquitin-proteosome system (UPS) in *S. cerevisiae* (Brandina et al. 2007). Prior studies have demonstrated the importance of UPS in protein quality control by eliminating proteins that are misfolded or aggregated (Kodroń et al. 2021). This suggests that finding tRNAs near genes related to the electron transport chain, specifically the UPS and cytochrome C, could indicate co-regulation of tRNAs and various aspects of protein control.

We also conducted species-specific ORA analysis for two species of interest: *S. cerevisiae* and *Y. lipolytica.* These two species were chosen because they have chromosome level genome assemblies, they have organism codes within the KEGG database, and they have recently published tRNA sequencing data. These two species also vary greatly in the total number of tRNA genes within their genomes: *S. cerevisiae* had 276, and *Y. lipolytica* had 511.

We found very little overlap between the two species and when comparing the overall data with the specific species. *S. cerevisiae* is used frequently in industry for the production of alcohol, and *S. cerevisiae* frequently utilizes rapid fermentation in aerobic and anaerobic conditions when glucose is present (Guzikowski et al. 2022). *Y. lipolytica* is used in industry due to its ability to store and utilize lipids (Beopoulos et al. 2009). These two yeast species are examples of the differences in the ecological roles of these yeast, and these differences could drive differences in genomic patterns. Another explanation could be that *S. cerevisiae* has around half the number of tRNA genes as *Y. lipolytica*, which could limit enrichment analysis. In both species, the genes near active tRNAs, as determined by tRNA sequencing, do not share functional categories with the genes found near all tRNAs. These differences are expected due to the differences in chromosome organization, as seen in Figure 1, and *Y. lipolytica* and *S. cerevisiae* have very different metabolic pathways. For example, *S. cerevisiae* uses fermentation, is Crabtree positive, and exhibits rapid cell growth (Guo et al. 2025). Conversely, *Y. lipolytica* is strictly aerobic and accumulates lipids (Bakkaiova et al. 2014).

**Figure 1:**
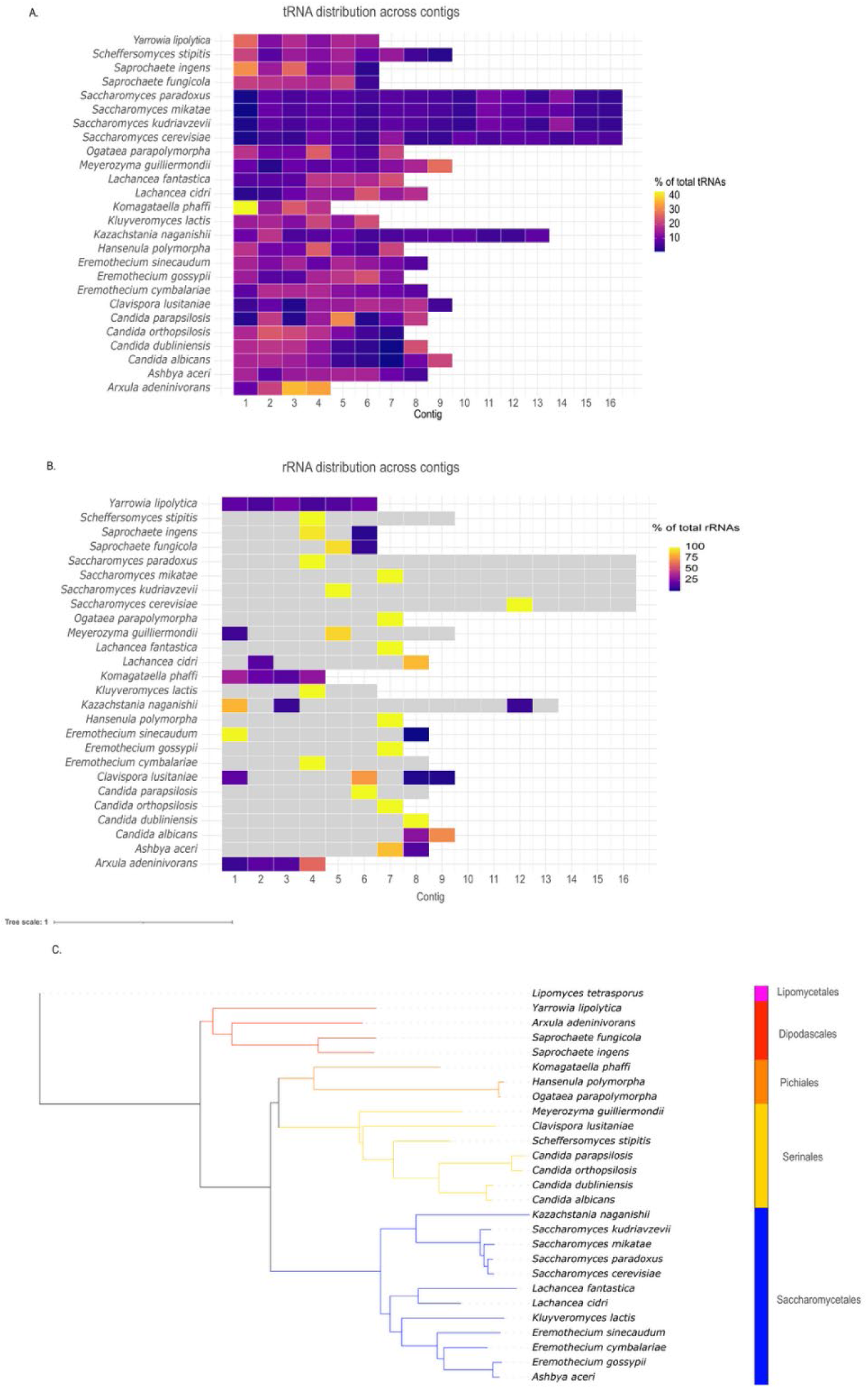
Chromosome level assemblies reveal tRNA and rRNA distribution is variable across species. (A) Predicted tRNAs were dispersed across chromosomes, except for some species such as *Komagataella phaffi* and *Arxula adeninivorans* which hadtRNAs more centralized on one or two chromosomes. (B) 13 out of 26 genomes had rRNAs predicted just on one chromosome. Conversely,*Yarrowia lipolytica* and *Arxula adeninivorans* hadrRNAs predicted on every one of their chromosomes. (C) Phylogenetic tree illustrating the 26 species subset. *Nakazawaeaw anatomiae* from the order Lipomycetales was used as an outgroup because it is present in the Saccharomycotina, but is absent from the 26 species subset.

Across 1,154 budding yeast genomes, we find that genes near tRNA are enriched in protein regulatory-related pathways and pathways utilizing electron transport. We hypothesize that the proximity of tRNA genes to ribosomal genes, in particular, leads to co-regulation of these translation-associated pathways. While yeast do not have neurons, yeast do have similar pathways in protein regulation, and that makes them a good model organism to study neurological disease (Ocampo and Barrientos 2011; Franssens et al. 2013; Mbefo et al. 2015; Zhao et al. 2018). Many of the KOs identified in this analysis in the neurodegenerative disease pathways were associated with proteasome, ion transport, and energy production. We also found that, despite broad patterns across all yeasts, individual genomes may differ in their genomic organization. Complete telomere-to-telomere genomes combined with tRNA-expression data will be needed to fully categorize tRNA genome organization.

### GO and GSEA Analysis of Nearby Genes

We also conducted GO analysis to support KEGG results. Of the 9,542 near-tRNA genes, 8052 had GO annotations associated with them. Both KEGG and GO showed high amounts of gene enrichment related to protein regulation, RNA processes, and ion transport. Shown in Figure 6A, RNA processing was the top result for GO enrichment for genes near tRNAs across the 1154 yeast genomes, and rRNA processing was the 5th highest result. rRNA is a key component of translation and is in line with the theme of gene proximity and the necessary components for translation being near each other. Other interesting results were TCA, which is involved in energy production, and the 4 different categories of ion transport. Ion transport was the second most prevalent category in the neurodegenerative category of KEGG ORA, and GO had results for proton transmembrane transport, inorganic cation transport, monoatomic cation transmembrane transport, and monoatomic ion transmembrane transport. These results support that genes near tRNA could be enriched in the necessary components for translation across both KEGG and GO analysis.

GSEA was also performed with KEGGcount as the ranking metric. This is further supported by the GSEA results, where protein export and RNA degradation were among the top results Figure 6B. mTOR, which plays a vital role in regulating protein synthesis and ribosome biogenesis, was particularly of interest as it was also a major result in ORA (Wang and Proud 2006). RNA degradation and protein export are intriguing GSEA results because they are also related to gene expression and properly folded proteins. Both regulate translation with RNA degradation, assessing the quality of mRNA and disposing of erroneous transcripts (Boehm et al. 2016), and protein export is involved in transporting newly synthesized proteins.

Evaluating genes near tRNAs with multiple metrics of enrichment analysis revealed similar patterns. Across the different annotation systems, GO, KEGG, and GSEA, there were consistent enrichment results.

### Distance from KEGG Pathways to tRNA

Translation-associated pathways were identified as enriched near tRNA genes. To further explore the two-dimensional organization of translation-associated genes and tRNAs, we expanded our analysis to all annotated genes in the target pathways. In this analysis, we compared the distance from every gene in a pathway or gene set to the nearest tRNA gene. The ribosomal pathways or genes included were: Ribosome pathway (map1234), ribosome protein (map3456), and Infernal RNA gene locations. Ribosome pathway (map1234) refers to genes that assist in ribosome assembly, but not necessarily the ribosome proteins themselves. Ribosomal proteins (map3456) refer to the genes encoding for the rRNAs, such as the large and small subunits. Proteasome was chosen due to protein processing results, and tRNA modification was chosen as another potential KEGG pathway that would be logical to be near RNAs. Glycolysis and Fatty Acid biosynthesis were controls that had a similar number of KOs as the other groups.

We found that translation-associated pathway genes were generally closer to tRNA genes than metabolic genes (Figure 2A). Of all the pathways tested, rRNA infernal had the highest density of distance values close to 0 (mean of −453bp and median of −4bp). The absolute values of these distances were also plotted as a density plot and overlaid (Figure 2B), and rRNA infernal still displays the highest density near 0. The metabolic pathways had a broader spread of distances than rRNA results and had a more even distribution around 0 instead of a large peak.

**Figure 2:**
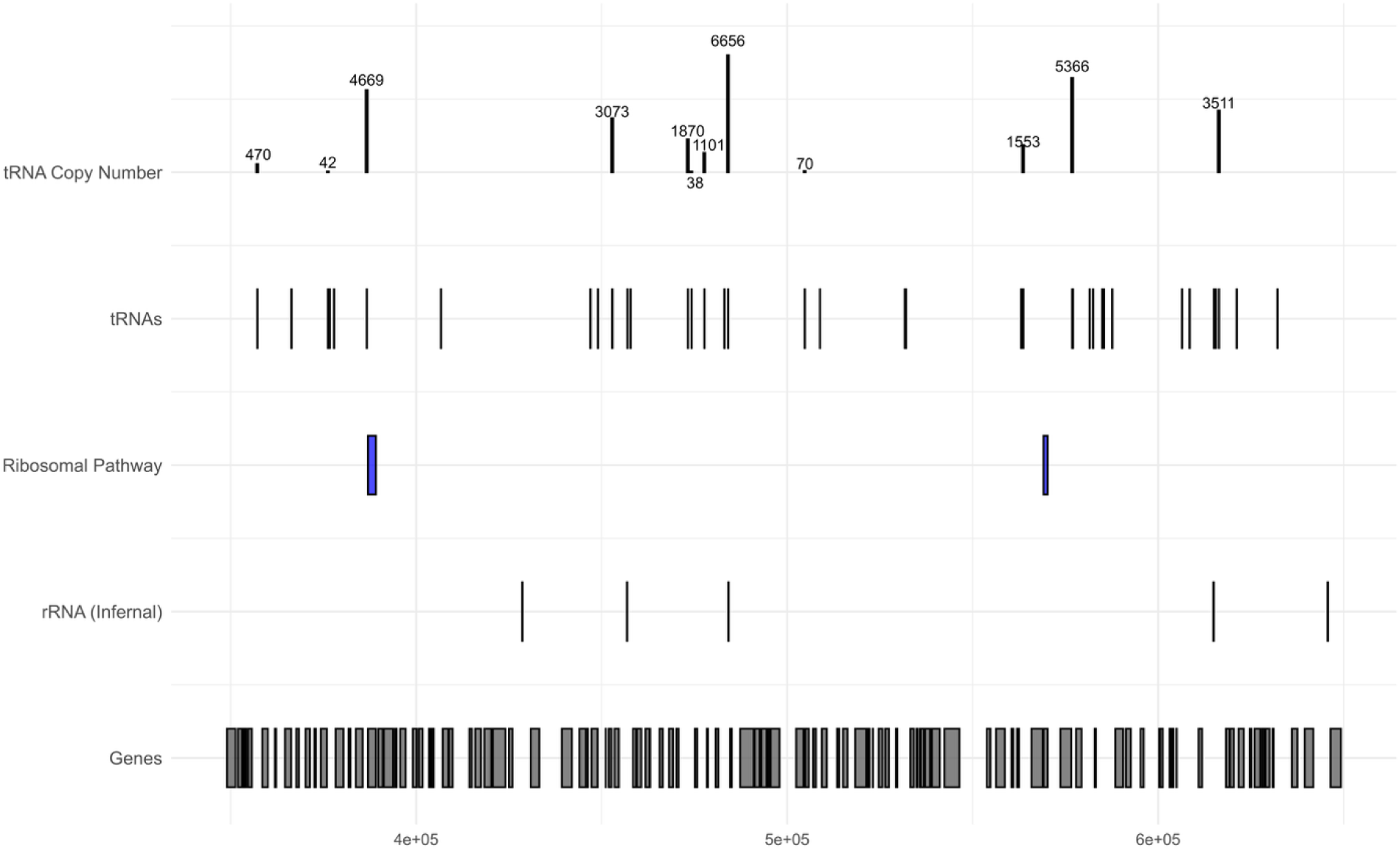
tRNA sequencing revealed a limited number of active tRNA genes in *Y. lipolytica*. Comparison of distances to ribosomal genes between active and inactive tRNAs genes revealed a representative region of the *Y. lipolytica* genome shows tRNAs with high copy number were near either the ribosomal proteins or the rRNA. tRNA copy number bars are scaled to allow for clearer visualization, and raw tRNA counts are listed on top of the copy number bars.

**Fig. 3:**
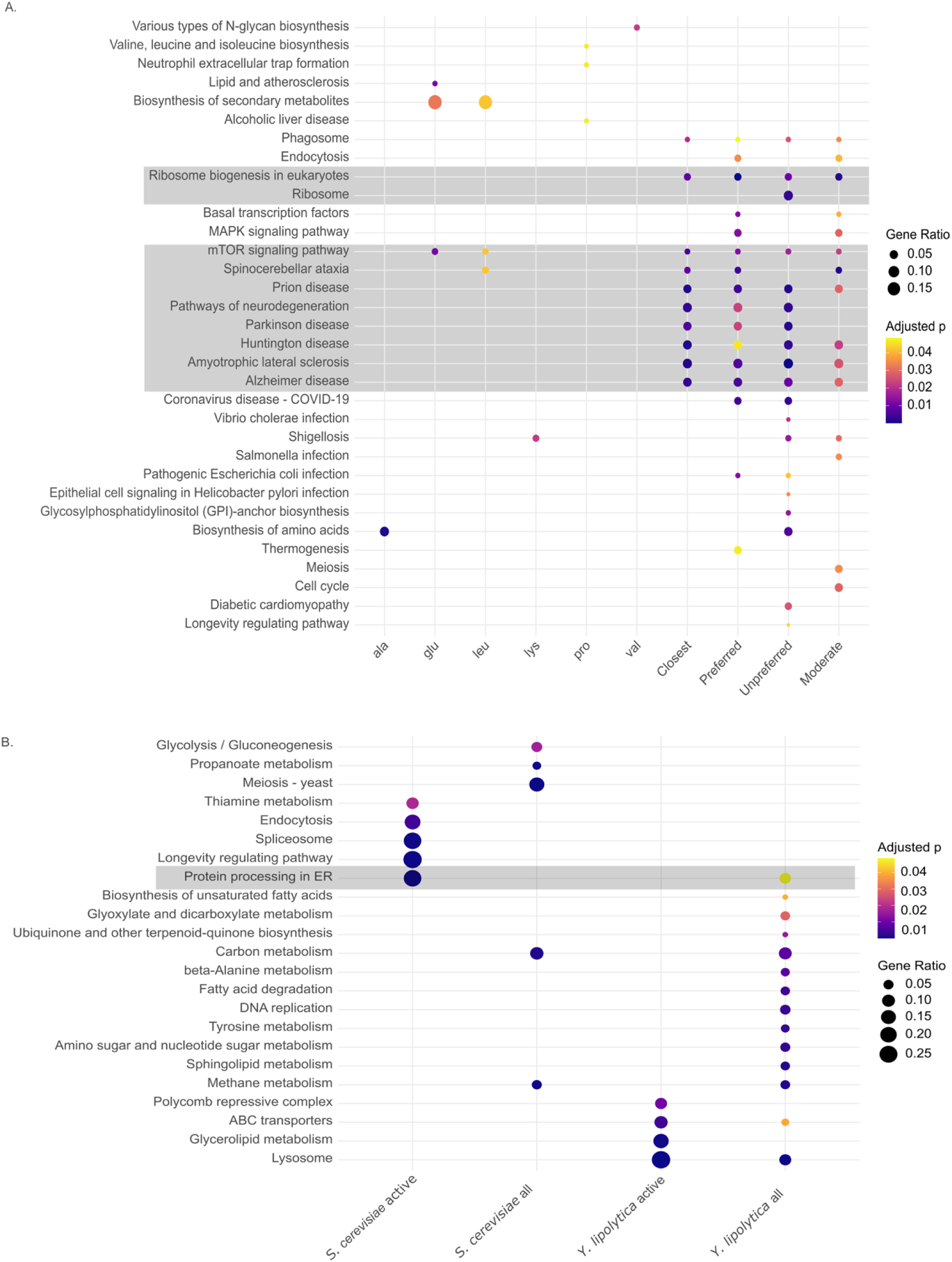
ORA revealed ribosome, neurodegenerative, protein, and metabolism KOs were overrepresented in genes near tRNAs. Bigger dots indicate higher gene ratio, and purple dots have a lower p-value. Gene ratio is the ratio of the number of genes part of a pathway over the total number of input genes. P-values were adjusted using the Benjamin-Hochberg (BH) multiple testing correction method. (A) There was consistent over-representation in pathways for the neurodegenerative disease and ribosome categories. The neurodegenerative diseases listed mostly have protein misfolding pathology, and most of the KEGG annotations for the neurodegenerative related pathways found in this study were related to proteasome, ion transport, and energy production. Groups included all 1154 yeast from the Saccharomycotina subphylum and were subset based on tRNA isotype, overall closest “Closest”, preferred (RSCU > 1.5), unpreferred (RSCU <0.5), and moderate (RSCU 0.5-1.5). (B) Species-specific analysis revealed metabolism pathways were overrepresented in genes near tRNAs. Enrichment patterns were different between the on genes and the window genes, with the only discernible pattern being a common theme of metabolism. “On genes” refers to genes near tRNAs that have a copy number greater than 12. “Window” is all the genes in a 2000 window of a tRNA.

**Fig. 5:**
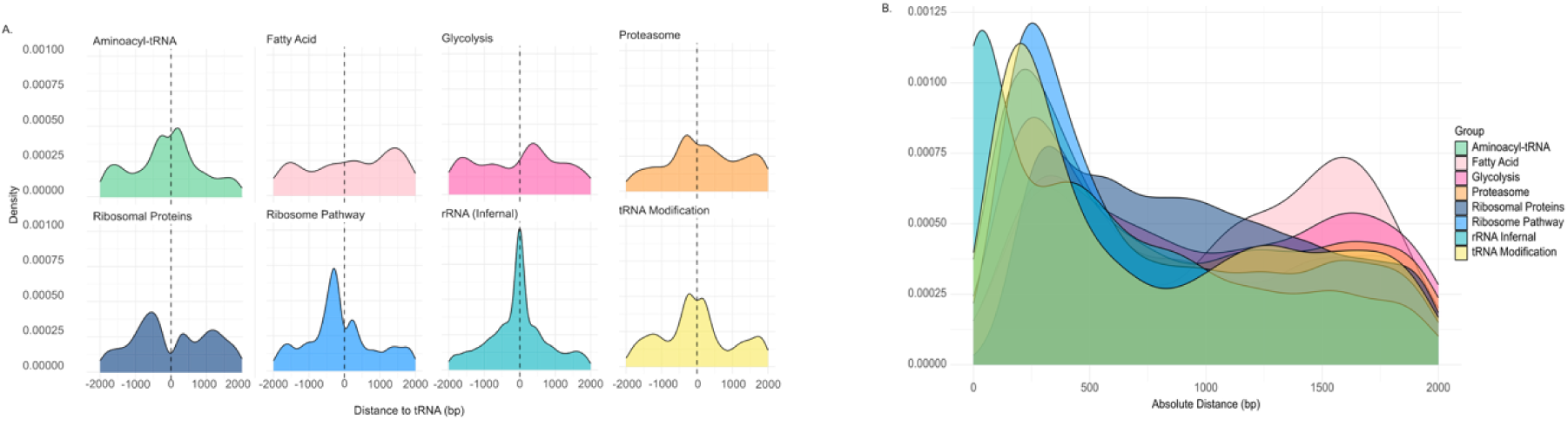
The distance from all genomic ribosome-associated genes to tRNAs was closer than expected based on comparison to selected metabolic gene categories. (A) rRNA genes had the highest density of values near the 0 distance from tRNA, and the KEGG ribosome pathway had the second highest density. The controls, fatty acid and glycolysis, did not show a peak near zero, suggesting they are not near tRNA genes. Density plots normalize the data so that the total area under the curve in the graph is 1. This allows for comparing distributions across multiple datasets. (B): rRNA genes showed a distribution of distances centralized around 0 with a qualitatively higher peak at 0 than the other groups. Sample sizes and total number of KEGG IDs is summarized in SB. All the groups are overlaid to provide a closer comparison.

**Fig. 6:**
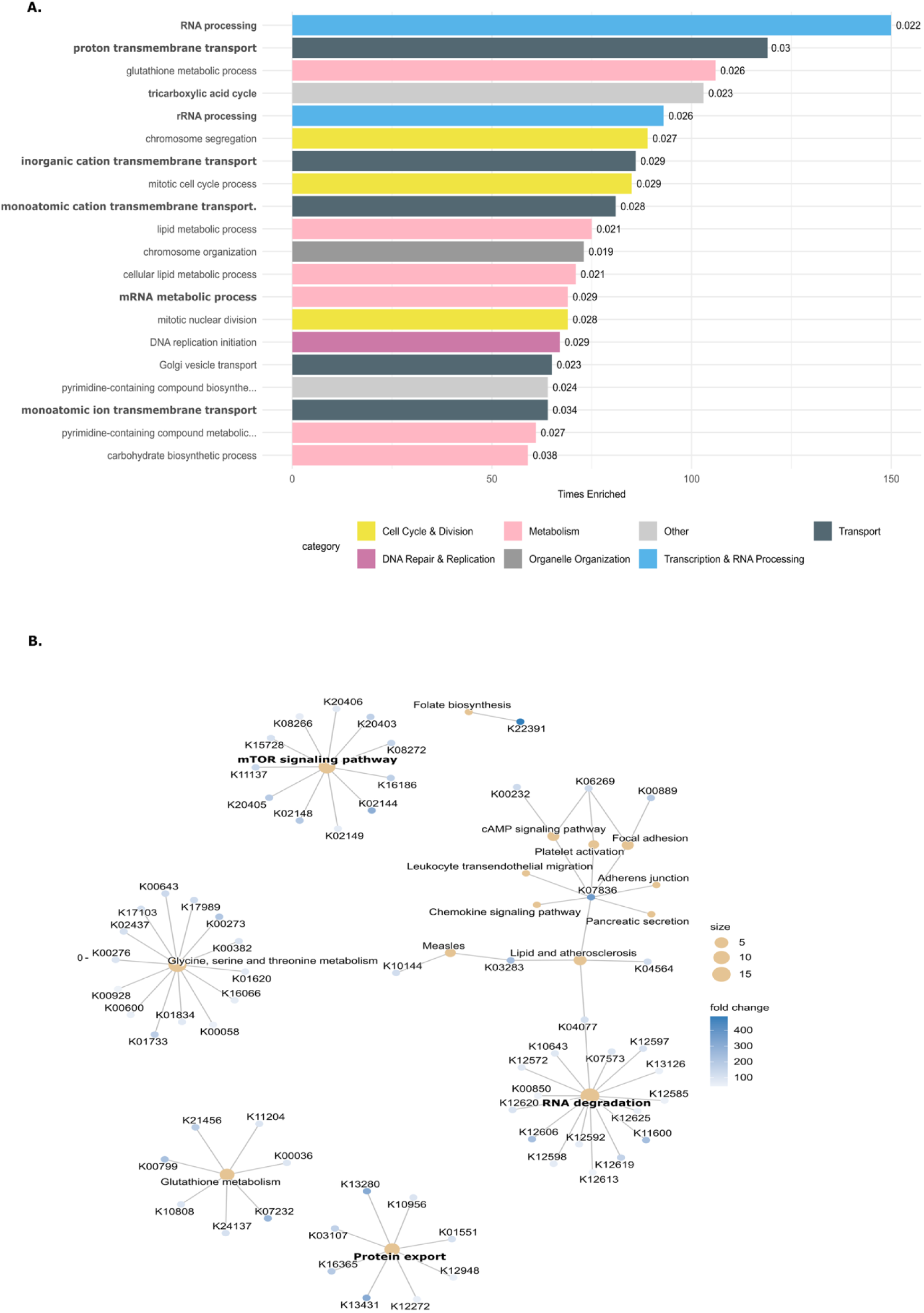
Gene Ontology and Gene Set Enrichment Analysis of genes near tRNA shows enrichment in genes important for translation and protein control. (A): GO RNA processing is a top result, and rRNA processing is one of the top terms as well. RNA processing is the parent GO term, which includes child terms of processing different types of RNA. This could be consistent with ribosomal genes from KEGG analysis. TCA cycle and the various categories of proton and ion transport are consistent with KEGG results that showed overrepresentation of genes enriched in processes of the electron transport chain. (B) CNET plot resulting from GSEA of genes near tRNAs. mTOR signaling pathway, protein export, and RNA degradation pathways from the GSEA analysis which are similar to the earlier in the ORA results of mTOR, ribosome biogenesis in eukaryotes, ribosome, and the various neurological hits that were part of protein quality control.

Statistical analysis of the pathway distance to tRNAs revealed that proteasome, ribosomal proteins, ribosomal pathway, rRNA infernal, and tRNA modification (Table 1) were significantly closer to tRNAs than the fatty acid biogenesis pathway, but only the ribosome pathway was significantly different from glycolysis. Examination of the statistics (Table 2) shows that the absolute value median and the mean of rRNA Infernal are closer to 0 than the other groups. This pathway analysis demonstrates that rRNAs are closer to tRNAs than other pathways.

**Table 1:**
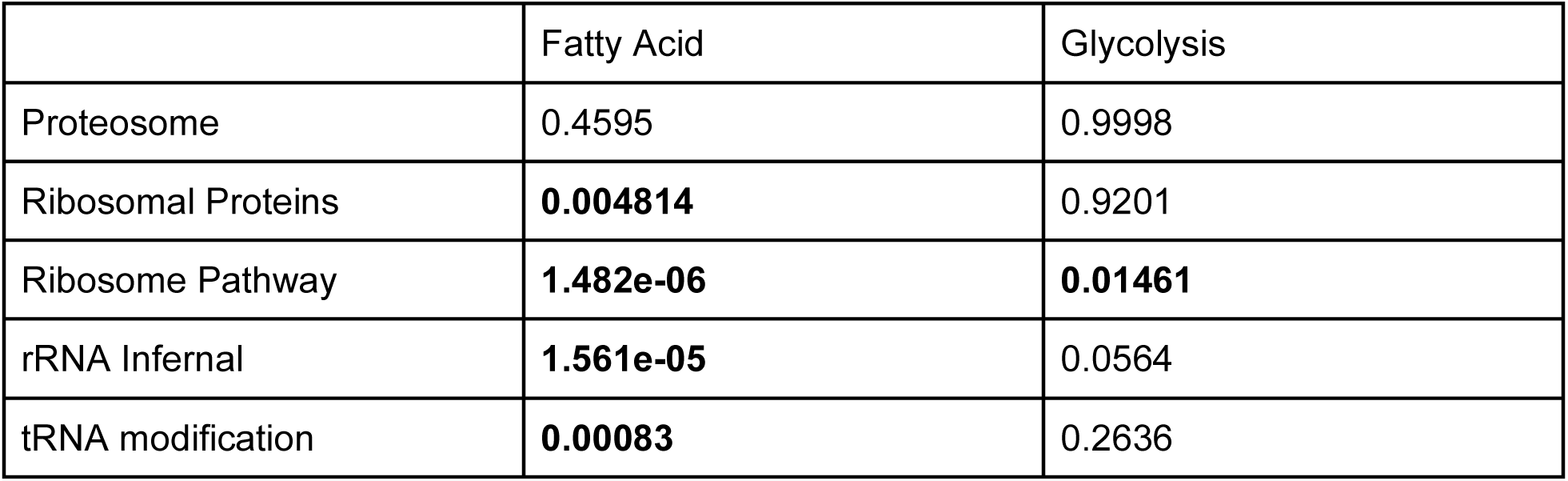
One-sided Wilcoxon Test: KEGG pathway (far left column) distances compared to Fatty Acid pathway and the glycolysis pathway to assess if the groups are significantly closer to tRNAs. The values reported in the table are the p-values from the One-Sided Wilcoxin Test. This table is answering the question of “Are the groups less than Fatty Acid and Glycolysis”?

|  | Fatty Acid | Glycolysis |
| --- | --- | --- |
| Proteosome | 0.4595 | 0.9998 |
| Ribosomal Proteins | **0.004814** | 0.9201 |
| Ribosome Pathway | **1.482e-06** | **0.01461** |
| rRNA Infernal | **1.561e-05** | 0.0564 |
| tRNA modification | **0.00083** | 0.2636 |

**Table 2:** Statistics for the absolute value of the distance of the KEGG pathways from tRNAs within 2000 base pairs. rRNA Infernal has the lowest median and mean of all the groups for distance from tRNAs.

|  | median | mean | n |
| --- | --- | --- | --- |
| rRNA Infernal | 458 | 614 | 3506 |
| Ribosomal Pathway | 564 | 778 | 4636 |
| Aminoacyl tRNA | 593 | 787 | 2303 |
| tRNA Modification | 624 | 806 | 3667 |
| Proteasome | 781 | 893 | 1930 |
| Ribosomal Proteins | 887 | 943 | 6262 |
| Glycolysis | 962 | 1004 | 713 |
| Fatty Acid | 1167 | 1062 | 395 |

## Conclusion

Current literature shows several steps from tRNA transcription, through processing to mature tRNA, but it still lacks the full story. Given the observations by others (Torres et al. 2019) and ourselves that tRNAs are differentially expressed across isotype and condition, there are likely other factors at play regulating tRNA. We demonstrated in this study that genomic organization could be another step in tRNA regulation. This was shown through consistent patterns of genes near tRNAs being enriched in various aspects of protein regulation across various computational methods, including ORA KEGG, GSEA KEGG, KEGG pathway, and GO analysis. Genes involved in the proteasome, TCA, and rRNA pathways were consistently enriched near tRNAs. These are all key components in protein production and regulation, and suggest that genes in close proximity to tRNAs are organized in such a way to encourage efficiency through intentional genomic organization.

One limitation of our work is that not all genes have annotated KEGG orthologs or GO IDs. Future studies should consider additional annotation methods to get a more robust overall view of patterns of gene enrichment of genes near tRNAs. Nevertheless, our dataset addresses these limitations through large scale analysis of 1154 genomes and integrating multiple tests that lead to similar conclusions. Additionally, our analysis also revealed between-species differences in tRNA and rRNA organization. Complete genomes would be needed for all known yeast species to conduct a thorough investigation of tRNA and rRNA evenness.

In conclusion, the analyses often lead to similar pathways, including RNA and protein regulation. mTOR, which regulates protein and ribosome biogenesis, was a significant result in both GSEA and ORA. RNA and rRNA were top GO results, which is in line with the KEGG pathway analysis that found that rRNA was closer to tRNA than other pathway groups. Numerous neurological diseases were found in KEGG ORA, but closer examination found that these KOs were largely proteasome and ion transport. Overall, these results demonstrate that genes near tRNAs are enriched in the components necessary for translation accuracy and efficiency.

## Acknowledgements

The authors thank the members of the LaBella Lab and the Y1000 Project (http://y1000plus.org) team.

## Funding Statement

This project was supported by the Lab LaBella within the Department of Bioinformatics and Genomics at UNC Charlotte. Computational analyses were conducted using the UNC Charlotte University Computing Resources. This work is supported by the NIH National Institute of General Medical Sciences (R34GM155455).

## Conflicts of Interest Statement

All authors declare no conflicts of interest.

## Data Availability

The data underlying this article are available in a Figshare repository that will be made available upon publication of the article.

## Notes

### Competing Interest Statement

The authors have declared no competing interest.

