## Supplemental Tables and Figures for "Genes near tRNAs are enriched in translational machinery"

Supplementary Tables & Figures – West et al 2026

Table S1: Number of chromosomes per species as reported by the literature and the number of contigs present in the assembly. Species that had chromosome number within 1 of the number of contigs were chosen for additional analysis.

| Species | Literature chromosomes | Literature source | Number of contigs |
|---|---|---|---|
| *Saprochaete fungicola* | 5 | https://journals.asm.org/doi/10.1128/mra.00092-19 | 6 |
| *Candida parapsilosis* | 8 | https://pmc.ncbi.nlm.nih.gov/articles/PMC6431126/ | 8 |
| *Clavispora lusitaniae* | 8 | https://pmc.ncbi.nlm.nih.gov/articles/PMC6935856/#:~:text=Five%20independent%20assemblies%20were%20obtained,and%20P4%20(see%20below). | 9 |
| *Meyerozyma guilliermondii* | 8 | https://pmc.ncbi.nlm.nih.gov/articles/PMC12720209/#:~:text=Data%20summary,1). | 9 |
| *Ashbya aceri* | 7 | https://academic.oup.com/g3journal/article/3/8/1225/6025766 | 8 |
| *Eremothecium gossypii* | 7 | https://www.ncbi.nlm.nih.gov/datasets/genome/GCA_000091025.4/ | 7 |
| *Eremothecium sinecaudum* | 8 | https://www.ncbi.nlm.nih.gov/datasets/genome/GCF_001548555.1/#:~:text=Table_title:%20Assembly%20statistics%20Table_content:%20header:%20%7C%20%7C,%7C%20RefSeq:%207%20%7C%20GenBank:%207%20%7C | 8 |
| *Yarrowia lipolytica* | 6 | https://www.mdpi.com/2309-608X/7/7/548 | 6 |
| *Kluyveromyces lactis* | 6 | https://www.ncbi.nlm.nih.gov/datasets/genome/GCA_000002515.1/#:~:text=Assembly%20methods,.labri.fr/Genolevures. | 6 |
| *Saccharomyces cerevisiae* | 16 | https://pmc.ncbi.nlm.nih.gov/articles/PMC3962479/#:~:text=Research%20regarding%20the%20genetics%20of,Oliver%2C%20England | 16 |
| *Kazachstania naganishii* | 12 | https://www.nature.com/articles/s42003-023-05285-0 | 13 |
| *Saprochaete ingens* | 5 | https://pmc.ncbi.nlm.nih.gov/articles/PMC6908801/ | 6 |
| *Candida dubliniensis* | 8 | https://www.kegg.jp/kegg-bin/show_organism?org=cdu | 8 |
| *Scheffersomyces stipitis* | 8 | https://www.kegg.jp/kegg-bin/show_organism?org=pic | 9 |
| *Candida albicans* | 8 | https://journals.asm.org/doi/10.1128/mbio.01205-18 | 9 |
| *Blastobotrys adeninivorans* | 4 | https://pmc.ncbi.nlm.nih.gov/articles/PMC4022394/ | 4 |

| | | | |
|---|---|---|---|
| *Saccharomyces paradoxus* | 16 | https://www.ncbi.nlm.nih.gov/datasets/genome/GCF_002079055.1/ | 16 |
| *Komagataella phaffii* | 4 | https://www.ncbi.nlm.nih.gov/datasets/genome/GCA_900235035.2/#:~:text=Download%20datasets%20API%20FTP,phaffii%20(Pichia%20pastoris)%20CBS7435. | 4 |
| *Ogataea parapolymorpha* | 7 | https://www.ncbi.nlm.nih.gov/datasets/genome/GCF_000187245.1/ | 7 |
| *Candida orthopsilosis* | 8 | https://pmc.ncbi.nlm.nih.gov/articles/PMC3338533/#:~:text=Our%20final%20C.,involving%20more%20than%20one%20gene. | 8 |
| *Ogataea polymorpha* | 7 | https://pmc.ncbi.nlm.nih.gov/articles/PMC9019687/#:~:text=The%20full%2Dlength%20genome%20of%20*Ogataea%20polymorpha*%20HU%2D11/CBS4732,structural%20variations%20between%20HU%2D11/CBS4732%20and%20NCYC495%20genomes | 7 |
| *Saccharomyces kudriavzevii* | 16 | https://www.ncbi.nlm.nih.gov/datasets/genome/GCF_947243775.1/#:~:text=Table_title:%20Assembly%20statistics%20Table_content:%20header:%20%7C%20%7C,%7C%20RefSeq:%2016%20%7C%20GenBank:%2016%20%7C | 16 |
| *Saccharomyces mikatae* | 16 | https://www.ncbi.nlm.nih.gov/datasets/genome/GCF_947241705.1/ | 16 |
| *Lachancea cidri* | 8 | https://journals.asm.org/doi/10.1128/msystems.01058-23#:~:text=Here%2C%20we%20used%20a%20combination,in%20the%20strain's%20phenotypic%20variation. | 8 |
| *Lachancea fantastica nom. nud.* | 7 or 8 | https://journals.plos.org/plosone/article?id=10.1371/journal.pone.0047834 or https://forge.inrae.fr/gryc-data/lachancea_fantastica-cbs_6924/-/tree/master?ref_type=heads | 7 |
| *Eremothecium cymbalariae* | 8 | https://pmc.ncbi.nlm.nih.gov/articles/PMC3276169/ | 8 |

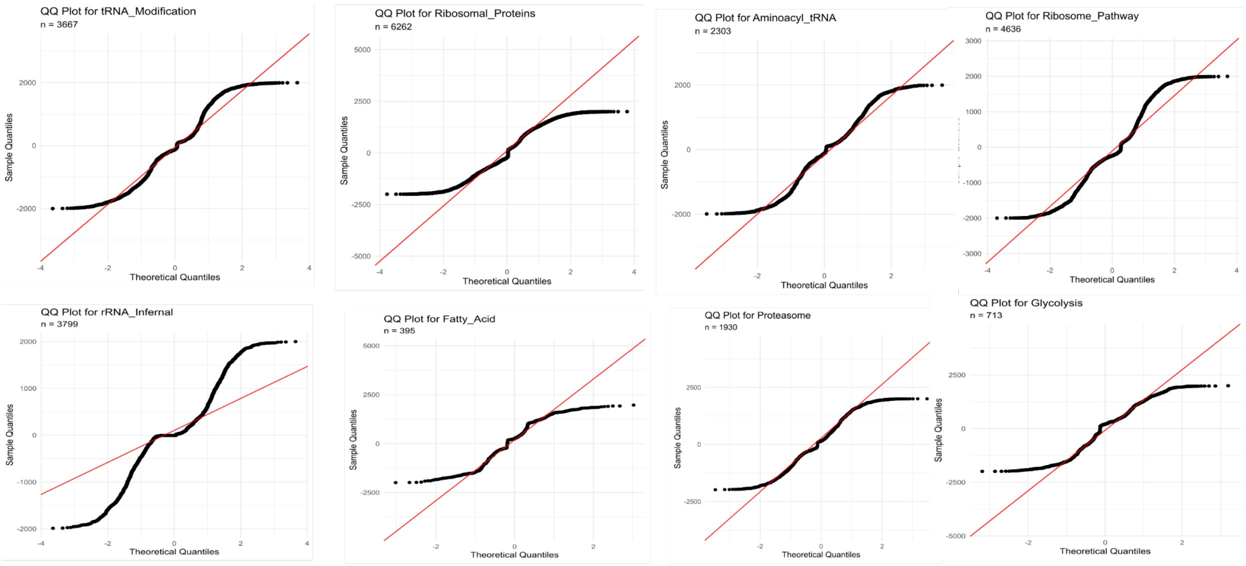


Figure S1: In order to determine which statistical test to use for tRNA modification, ribosomal protein, aminoacyl tRNA, ribosome pathway, rRNA infernal, fatty acid, glycolysis, and proteasome groups distance from tRNAs, qqplots were generated. None of the qqplots showed a normal distribution. A non-normal distribution requires a non-parametric test, so the results of the qqplots indicate the need for a Wilcoxon rank-sum test.

Table S2: The different groups distance from tRNA was tested for normality with the Shapiro Wilks and Anderson Darling tests. These tests reveal that the data is not normally distributed, and is in agreement with the non-normality results from the qqplots. Therefore a Wilcoxin rank-sum test will be used for statistical testing.

| Group | n | shapiro_p | ad_p | shapiro_normal | ad_normal |
|---|---|---|---|---|---|
| **Aminoacyl_tRNA** | 2303 | 5.49498527327931E-07 | 3.7E-24 | No | No |
| **Fatty_Acid** | 395 | 5.53315769272589E-13 | 8.2400835445722E-21 | No | No |
| **Glycolysis** | 713 | 9.67753981224753E-13 | 2.06627507867344E-23 | No | No |
| **Proteasome** | 1930 | 1.10759053981797E-09 | 3.7E-24 | No | No |
| **Ribosomal_Proteins** | 6262 | 1.36239446986738E-10 | 3.7E-24 | No | No |
| **Ribosome_Pathway** | 4636 | 3.49617191642406E-10 | 3.7E-24 | No | No |
| **rRNA_Infernal** | 3799 | 4.761939513843E-13 | 3.7E-24 | No | No |
| **tRNA_Modification** | 3667 | 5.30593275913337E-09 | 3.7E-24 | No | No |

Table S3: KEGGs related to neurological function were enriched near tRNAs. Since yeast do not have a neurological system, the KEGG IDs were further investigated and grouped into plausible pathways. This table explains the grouping methodology.

<table>
<tr><th>Group</th><th>Total_Count</th><th>Num_KEGGs</th><th>REGEX</th></tr>
<tr><td>Ion Transport</td><td>320</td><td>55</td><td>transporter|channel|P-type.*|VDAC|voltage-dependent anion channel|solute carrier|NADH dehydrogenase|F-type H\\+-transporting ATPase|succinate dehydrogenase</td></tr>
<tr><td>Proteasome</td><td>217</td><td>31</td><td>Proteasome</td></tr>
<tr><td>Mitochondrial / Redox</td><td>16</td><td>7</td><td>cytochrome|ubiquinol|oxidase|superoxide|catalase|glutathione|mitofusin|OMA1</td></tr>
<tr><td>Signaling / Kinases</td><td>39</td><td>11</td><td>kinase|MAPK|mTOR|phosphatase</td></tr>
<tr><td>Transcription / Nuclear transport</td><td>33</td><td>32</td><td>RNA polymerase|transcription|TFIID|histone|nuclear|nucleoporin|export</td></tr>
<tr><td>Autophagy</td><td>30</td><td>6</td><td>autophagy|beclin|ULK|ATG</td></tr>
<tr><td>Cytoskeleton</td><td>21</td><td>4</td><td>Tubulin|kinesin|dynactin</td></tr>
<tr><td>Ubiquitin system</td><td>20</td><td>9</td><td>ubiquitin|E1|E2|NEDD8</td></tr>
<tr><td>Other</td><td>15</td><td>13</td><td></td></tr>
<tr><td>Other signaling / Regulatory</td><td>11</td><td>5</td><td>Ras|calmodulin|thioredoxin</td></tr>
<tr><td>Chaperones / ER</td><td>9</td><td>5</td><td>BiP|stress-induced|peptidyl-prolyl|heat shock|ATP-dependent RNA/DNA helicase</td></tr>
<tr><td>Vesicle / Transport / Trafficking</td><td>7</td><td>6</td><td>clathrin|AP-2|TOM40|SEC13</td></tr>
<tr><td>Translation / Ribosome</td><td>5</td><td>1</td><td>translation initiation factor|ribosomal|ubiquitin-large subunit ribosomal</td></tr>
<tr><td>Total</td><td>743</td><td>185</td><td></td></tr>
</table>
